# Allelic polymorphism at foxo contributes to local adaptation in Drosophila melanogaster

**DOI:** 10.1101/471565

**Authors:** Nicolas J. Betancourt, Subhash Rajpurohit, Esra Durmaz, Daniel K. Fabian, Martin Kapun, Thomas Flatt, Paul Schmidt

**Affiliations:** Department of Biology, University of Pennsylvania, Philadelphia, USA; Ahmedabad University, Division of Biological and Life Sciences, Ahmedabad, India; Department of Ecology and Evolution, University of Lausanne, Lausanne, Switzerland; Department of Biology, University of Fribourg, Fribourg, Switzerland; Department of Genetics, University of Cambridge, Cambridge, United Kingdom; European Bioinformatics Institute (EMBL-EBI), Hinxton, United Kingdom

**Keywords:** *foxo*, cline, size, starvation tolerance, genetic architecture

## Abstract

The insulin insulin-like growth factor signaling pathway has been hypothesized as a major determinant of life history profiles that vary adaptively in natural populations. In *Drosophila melanogaster*, multiple components of this pathway vary predictably with latitude; this includes *foxo*, a conserved gene that regulates insulin signaling and has pleiotropic effects on a variety of fitness-associated traits. We hypothesized that allelic variation at *foxo* underlies genetic variance for traits that vary with latitude and reflect local adaptation. To evaluate this, we generated recombinant outbred populations in which the focal *foxo* allele was homozygous and fixed for either the allele common at high latitude or low latitude and the genomic background was randomized across 20 inbred lines. After eight generations of recombination, experimental populations were phenotyped for a series of traits related to gene function. Our results demonstrate that natural allelic variation at *foxo* has major and predictable effects on body size and starvation tolerance, but not on development time. These patterns mirror those observed in natural populations collected across the latitudinal gradient in the eastern U.S.: clines were observed for starvation tolerance and body size, but development time exhibited no association with latitude. Furthermore, differences in size between *foxo* genotypes were equivalent to those observed between populations sampled from the latitudinal extremes, although contribution to the genetic variance for starvation tolerance was less pronounced. These results suggest that allelic variation at *foxo* is a major contributor to adaptive patterns of life history variation in natural populations of this genetic model.

## 1. INTRODUCTION

Elucidating the molecular, mechanistic basis of adaptive differentiation for complex traits in natural populations remains a fundamental goal in evolutionary biology. Outlining the genotype-phenotype map and subsequent investigation into mechanism requires, to some extent, identification of the genes and variants that underlie genetic variance realized in the wild. However, fitness traits are often complex, with a highly polygenic architecture (e.g., Arnegard et al. 2014; McCown et al. 2014; Savolainen et al. 2013). The likelihood of effectively mapping complex traits to causative polymorphism varies between two classical viewpoints regarding architecture for quantitative traits: many loci of small effect vs. few loci of large effect (Rockman 2012; Wellenreuther and Hansson 2016; Roff 2017; Boyle et al. 2017; Barton et al. 2017). While these perspectives may be artificially polarized, many empirical advances in understanding the mechanistic basis of adaptation in sexual, outbred populations include both a clear identification of traits that drive local adaptation as well as an apparently simple genetic architecture, with at least one locus of major effect (e.g., Colosimo et al. 2005; Comeault et al. 2015; van’t Hof et al. 2016; Lamichhaney et al. 2016; Jones et al. 2018). In such examples, alleles segregating at a locus of major effect can then be examined for functional differences that affect performance and fitness (e.g., Laurie and Stam 1988; Manceau et al. 2011; Cheviron et al. 2012; Chakraborty and Fry 2016). What is unclear, however, is whether alleles at loci underlying local adaptation are typically of large effect, or whether the effect size of such naturally occurring variants is effectively negligible for any molecular or functional investigation (e.g., Lewontin 1974; Rockman 2012).

Body size is a trait commonly associated with fitness (Brown et al. 1993; Blanckenhorn 2000; Bonnet et al. 2017). In a variety of taxa, size also varies predictably across environmental gradients such as those associated with latitude (James et al. 1995; Huey et al. 2000; Ashton 2002; Blanckenhorn and Demont 2004; Stillwell et al. 2007); such clines suggest that body size is affected by spatially varying selection and contributes to local adaptation (Partridge and Coyne 1997; Stillwell 2010). While size-related traits are in general highly polygenic (e.g., Boyle et al. 2017), major effect loci have also been observed (e.g., Sutter et al. 2007). In particular, multiple components of the insulin insulin-like growth factor signaling pathway (IIS) can regulate size (e.g., Colombani et al. 2005; Sutter et al. 2007); in particular, the forkhead box-O transcription factor gene *foxo* is a major regulator of IIS and impacts size as well as a variety of other traits associated with fitness (Libina, Berman, & Kenyon, 2003; Kramer, Davidge, Lockyer, & Staveley, 2003; Kramer, Slade, & Staveley, 2008; Hwangbo, Gersham, Tu, Palmer, & Tatar, 2004; Fielenbach & Antebi, 2008; Mattila, Bremer, Ahonen, Kostiainen, & Puig, 2009). Thus, the analysis of variation in body size offers an excellent system in which to examine aspects of genetic architecture and the integration between allelic and phenotypic patterns in natural populations.

In *Drosophila melanogaster*, body size increases with increasing latitude on multiple continents (Coyne and Beecham 1987; James et al. 1995, 1997; Karan et al. 1998). Such latitudinal patterns are mirrored by altitudinal clines where size increases with increasing altitude (Fabian et al. 2015; Lack et al. 2016). These parallel and replicated patterns suggest that patterns of size variation are adaptive and directly associated with thermally mediated selection (reviewed in Stillwell 2010). However, it remains unknown why small body size is associated with high fitness at low latitudes and large size with high fitness at high latitudes. De Jong and Bochdonavits (2003) hypothesized that adaptive patterns of size variation are driven primarily by one or more components of the IIS pathway; the simple prediction is that any causative variants would also exhibit pronounced and replicated allele frequency clines. Analysis of PoolSeq data has shown that multiple IIS genes (e.g., *Pi3K*, *foxo*, *InR*) are segregating for many alleles that vary predictably with latitude in *D. melanogaster* (Kolaczkowski et al. 2011; Fabian et al. 2012; Bergland et al. 2014; Kapun et al. 2016; Machado et al. 2018). The question is whether these clinal alleles are distinct with respect to gene function and, at least in part, underlie the observed patterns of local adaptation in size (Paaby et al. 2014).

Here, we examine the functional significance of naturally occurring allelic variation at the forkhead box-O transcription factor gene *foxo* and evaluate whether the effects of allelic variation are consistent with patterns of variation among natural populations. We use a specific *foxo* allele, previously identified in PoolSeq data as being strongly clinal in the eastern U.S. (Fabian et al. 2012), as a marker for candidate alleles of functional significance. If size in *D. melanogaster* is highly polygenic with no loci of major effect, then alleles segregating at *foxo* should be effectively and functionally equivalent in laboratory- or field-based functional assays. However, if *foxo* is a major locus for body size in this species, then effect size may be of sufficient magnitude to be detectable. If allelic variation at *foxo* substantially contributes to phenotypic variation and local adaptation, then the effects of *foxo* alleles on phenotype should also be concordant with patterns observed in natural populations. Such patterns should also be trait-specific and restricted to those related to gene function.

## 2. MATERIAL AND METHODS

### 2.1 Identification of SNPs/alleles that vary predictably with latitude

Fabian et al. (2012) identified a series of SNPs in *foxo* that exhibited high *F_ST_* in pooled sequencing of natural populations derived from Florida (low latitude), Pennsylvania (mid latitude), and Maine, U.S.A. (high latitude). From this analysis, we identified a candidate *foxo* allele based on the nucleotide state at two SNPs spanning approximately 2kb. Further details are provided in Durmaz et al. (2018).

To further examine clinal patterns associated with the focal low- and high-latitude *foxo* haplotypes, we analyzed ten populations collected along the latitudinal gradient of the U.S. east coast and sequenced as pools (Bergland et al. 2014; Kapun et al. 2016; Machado et al. 2018). We restricted our analyses to high-confidence SNPs that were polymorphic in the *Drosophila* Genetic Reference Panel (DGRP) dataset (Mackay et al. 2012; Langley et al. 2012) and isolated a total of 1372 SNPs located inside or within 2kbp up- and downstream of the annotated *foxo* gene. To provide a null context for allele frequency differentiation, we isolated 20,000 SNPs located in short introns (<60bp) and at least 100 kbp distance from the breakpoints of common cosmopolitan inversions (Corbett-Detig et al. 2012) that are consistent with patterns of evolutionary neutrality (Parsch et al. 2010; Clemente & Vogl 2012). For each of these neutral and *foxo*-associated SNPs, we tested for significant correlations between latitude and allele frequencies using generalized linear models (GLM) with a binomial error structure of the form: *y*i = *L* + *ε*i, where *y*i is the allele frequency of the *i*^th^ SNP, *L* is the continuous factor “Latitude” and *ε*i is the binomial error of the *i*^th^ SNP. We further assessed whether allele frequency changes of the two candidate SNPs that constitute our focal haplotype were more clinal than neutral SNPs or other SNPs located within or in the proximity of *foxo*. To this end, we compared the -log_10_(*p*)-values from the GLMs for each of the two focal SNPS to distributions of -log_10_(*p*)-values from GLMs of either neutral SNPs or non-candidate SNPs associated with *foxo*. We subsequently calculated empirical cumulative distribution functions (ECDF; with total area =1) based on -log_10_(p)-values from all neutral or non-candidate *foxo* SNPs in *R* (R Development Core Team 2009). To test if the significance values associated with the two focal SNPs were greater than the 95 percentiles of each significance distribution, we integrated over the area under each ECDF with values larger than the significance of each candidate SNP (see gray areas limited by red dashed lines in Figure 1) and subtracted the integral value from 1, which represents the total area of the ECDF.

**Figure 1.**
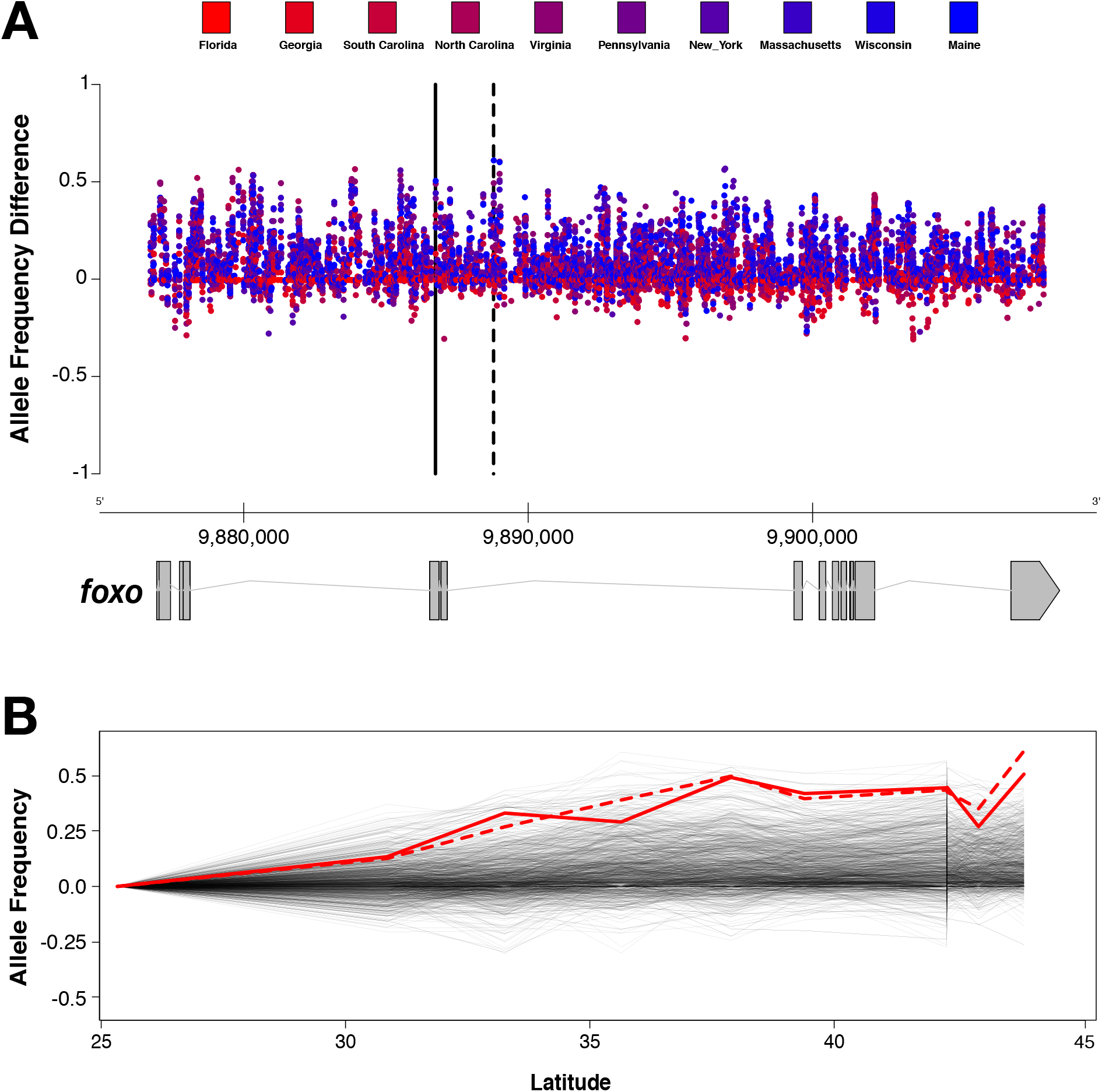
Allele frequency changes for *foxo*-associated SNPs in 10 populations sampled along the North American east coast. Both plots show allele frequency differences conditioned to increase from south to north, with frequencies in Florida being set to zero. Panel A shows allele frequency changes for all SNPs according to their genomic position. Here, the *foxo* candidate SNPs are highlighted by two vertical black lines (solid: 3R: 9,892,517; dashed: 3R: 9,894,449; *Drosophila melanogaster* reference genome v6). Panel B shows how allele frequencies change with latitude. The two *foxo* candidate SNPs are highlighted in red (solid: 3R: 9,892,517; dashed: 3R: 9,894,449).

To investigate and visualize the relative patterns of allele frequency change for the two focal SNPs with respect to all other SNPs located inside or in close proximity to *foxo*, we conditioned the alleles with lower frequencies in Florida compared to the population in Maine for each *foxo*-associated SNP. Allele frequencies in Florida were set to zero and we then calculated the allele frequency differences relative to Florida for all other populations.

### 2.2 Constructing recombinant population cages

Based on the combination of the results of Fabian et al. (2012) and the DrosRTEC (*Drosophila* Real-Time Evolution Consortium) sequencing effort (Machado et al. 2018), we identified individual lines in the DGRP (Mackay et. al 2012) that were homozygous for the candidate *foxo* allele that was at high frequency in high-latitude populations (hereafter, the high-latitude allele), and lines that were fixed for the *foxo* allele that was at high frequency in low-latitude samples (hereafter, the low-latitude allele). Scripts for this filtering of the DGRP based on nucleotide state and locus are provided in the raw data (Data Dryad submission XXXXX). Two biological replicate population cages, denoted as sets A and B, were established using 20 independent lines per cage per *foxo* allele; thus, each cage was constructed with a completely distinct and independent set of inbred lines. Each population cage was founded using 10 individuals of each sex, from age- and density-controlled cohorts, from each of the 20 inbred DGRP founding lines.

Biological replicates were further split into two experimental replicates in the first generation of founding and cultured independently thereafter. After establishment, each population cage was cultured at a population size of ~2000 adults on standard cornmeal-molasses medium for 8 discrete generations of outcrossing under conditions of constant temperature (25°C) and photoperiod (12L:12D) in Percival I36VL incubators. Thus, at the end of the experimental period, we generated replicate population cages in which the focal *foxo* allele was homozygous and fixed for either the high-latitude or low-latitude allele and the genomic background was randomized across the 20 inbred lines used to found each respective cage (Paaby et al. 2014; Behrman et al. 2018). To further test for genome-wide patterns of genetic differentiation among the recombinant outbred populations (ROP) fixed for either the high or low-latitude haplotypes, we calculated SNP-wise *F*_ST_ based on the method of Weir and Cockerham (1984) for sets A and B separately using custom software. After the eight generations of density-standardized culture under standardized environmental conditions, we established replicate density-controlled vial cultures (30 ± 10 eggs/vial) for subsequent phenotyping of the high-latitude and low-latitude *foxo* alleles.

### 2.3 Isofemale lines from natural populations

Thirty isofemale lines were randomly selected from each of six previously collected outbred populations along the east coast of the U.S. to serve as a latitudinal comparison to the recombinant *foxo* populations (described in Rajpurohit et al. 2017, 2018: Homestead, FL (HFL), Jacksonville, FL (JFL), Charlottesville, VA (CVA), Media, PA (MPA), Lancaster, MA (LMA), and Bowdoin, ME (BME)). The individual lines from each population were maintained on a standard 21d culture regime under the same environmental conditions as the *foxo* recombinant cages. Prior to phenotyping, each isofemale line was cultured for two generations at low density (30±10 eggs/vial) at 25°C, 12L:12D; in the third generation, freshly eclosed flies were collected in daily cohorts and used in the phenotypic assays described below.

### 2.4 Phenotype assays

In all assays, *foxo* recombinant cages were tested simultaneously at three independent timepoints and the data partitioned into blocks. For assessment of starvation resistance, virus infection of the cages precluded running three independent blocks, and a single timepoint was included in the analysis. For the natural populations, all lines from all populations were assayed simultaneously for all phenotypes using discrete 1d cohorts for each phenotype.

*Starvation resistance*. For the *foxo* recombinant outbred populations, embryos were collected from each cage in two replicate glass culture bottles and density was standardized at 150±10 eggs per bottle. Isofemale lines from the natural populations were transferred into replicate vials and density standardized at 30±10 embryos per vial. All culture was done under standard conditions of 25°C and a photoperiod of 12L:12D. Upon eclosion, mixed sex daily cohorts were collected over 3d and subsequently aged to 5d. The flies were then separated by sex into replicate groups of 10 and placed into glass vials equipped with a small cotton ball saturated with 1 mL of water. Samples were placed in an incubator at 25°C, 12L:12D and mortality was recorded at four timepoints per day (9AM, 1PM, 5PM, and 9PM) until all flies had died. The *foxo* cages were assayed in three replicates per cage population; all isofemale lines from each of the natural populations were also assayed.

*Development time*. For the *foxo* recombinant outbred populations, eggs were collected from each cage over a 3h window using large Petri dishes containing standard medium supplemented with live yeast. The collected eggs were then counted and distributed in groups of 30 into three replicate vials per cage. For the natural populations, embryos from all isofemale lines were similarly collected over 3h in small collection receptacles. Embryos were counted and distributed to new collection vials, with density also standardized at 30 embryos per vial. All experimental material was subsequently cultured as previously described in Percival I36VL incubators with constant and standardized humidity, temperature, and photoperiod. All experimental vials were checked four times daily (9AM, 1PM, 5PM, 9PM); eclosion events and sex were recorded.

*Body size/morphology*. For both the *foxo* cage as well as the natural populations, flies from the development assay were transferred to new vials, allowed to mate and age for 5d post eclosion, then were preserved in 95% ethanol in eppendorf tubes for subsequent size measurements. From these preserved samples, 10 flies for each sex were randomly sampled and measured from each *foxo* ROP cage and 5 flies for each sex were measured for each isofemale line from the natural populations. Body size measurements (thorax length and wing area) were recorded using a Leica MZ9.5 microscope mounted with an Olympus DP73 camera with CellSens standard measuring software. Thorax length was measured as the longest length across the dorsal shield; wing area was defined as a polygon using a standardized series of veinous landmarks. The ratio of total wing area to thorax length, indicative of wing loading (Azevedo et al. 1998; Gilchrist et al. 2000), was also calculated and subsequently analyzed.

### 2.5 Statistical analysis

For the natural populations, data were analyzed separately by sex. Isofemale line was considered a random variable and all data were analyzed using a restricted maximum likelihood ANOVA with population as a fixed effect. For all other traits other than starvation tolerance, experimental block (N=3) was also included as a covariate. For the *foxo* experimental population cages, a similar nested ANOVA was run independently for both sexes in which *foxo* allele, biological replicate (set), and experimental replicate (cage) were included as predictors.

## 3. RESULTS

### 3.1 DrosRTEC Pooled Population Genome Sequencing

By analyzing extensive genome-wide Pool-Seq data from ten populations sampled along the U.S. east coast generated by the DrosRTEC consortium (Bergland et al. 2014; Kapun et al. 2016; Machado et al. 2018), we show that numerous SNPs associated with *foxo* exhibit steep latitudinal clines and extensive differentiation as a function of geography (Figure 1). Notably and consistent with previous observations by Fabian et al. (2012), the two focal *foxo* candidate SNPs that are highlighted by black and red lines in Figures 1A and 1B, respectively, exhibit strong allele frequency change between the populations at lowest and highest latitudes. In fact, clinal patterns of the two candidate SNPs were more pronounced than 95% of neutrally evolving SNPs located in short introns (>95.94% for 3R:9,892,517 and >98% for 3R:9,894,559, respectively; Figure 2 A, B) and 91% of all non-candidate SNPs located within or close to *foxo* (>91.22% for 3R:9,892,517 and >96% for 3R:9,894,559, respectively; Figure 2 C, D).

**Figure 2.**
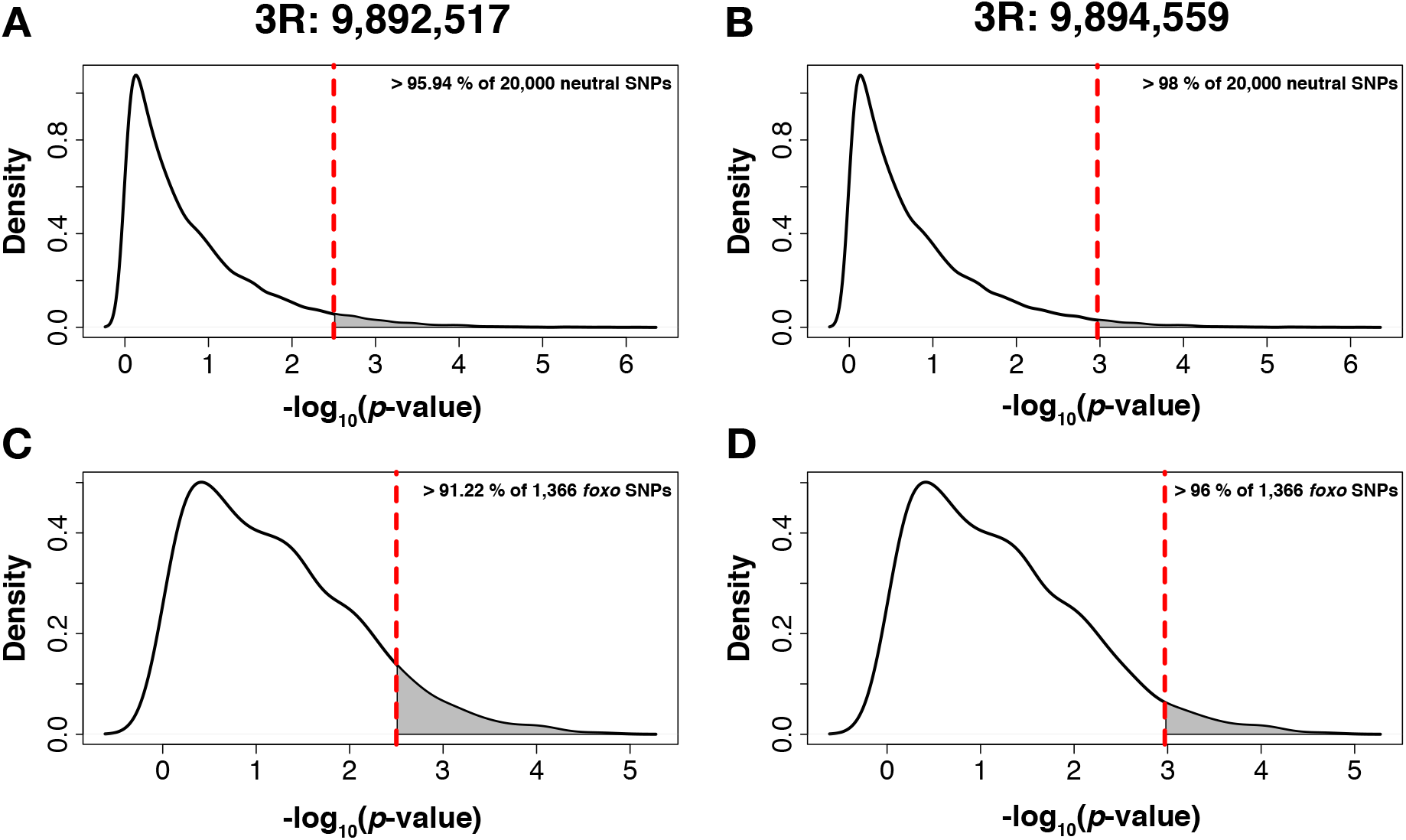
Empirical cumulative density functions (ECDF; total area = 1) calculated from the distribution of -log_10_(*p*-values) for generalized linear models that test for associations between allele frequencies and latitude in 20,000 neutrally evolving SNPs (Panels A, B) and 1,372 non-candidate SNPs located inside or within 2 kbp distance to *foxo*. The vertical dashed lines indicate the significance values of the two candidate SNPs 3R: 9,892,517 (Panels A, C) and 3R: 9,894,449 (Panels B, D). The grey areas limited by the dashed line indicate the percentiles of neutral or noncandidate *foxo* SNPs with significance values larger than the candidates.

In establishing the recombinant outbred populations (ROPs) using the DGRP panel of inbred lines, the goal was to use candidate SNPs as markers of functional effects for naturally occurring alleles or haplotypes at this locus (Paaby et al. 2014; Berhman et al. 2018). The utility of this method is predicated on using a sufficient number of independent inbred lines such that no other position in the genome, other that the candidate site(s), is fixed or highly differentiated between experimental sets. In Figure 3, *F_ST_* between experimental cages (*foxo* allele AT vs. *foxo* allele GG) is plotted as a function of chromosomal position for all SNPs segregating in the biological replicate sets A (Figure 3A) and B (Figure 3B). While there are multiple sites on each chromosome arm with *F_ST_* > 0.4 between the sets of lines used to construct the alternative *foxo* allelic cages, only the candidate sites are fixed between the cages that comprise the allelic states.

**Figure 3.**
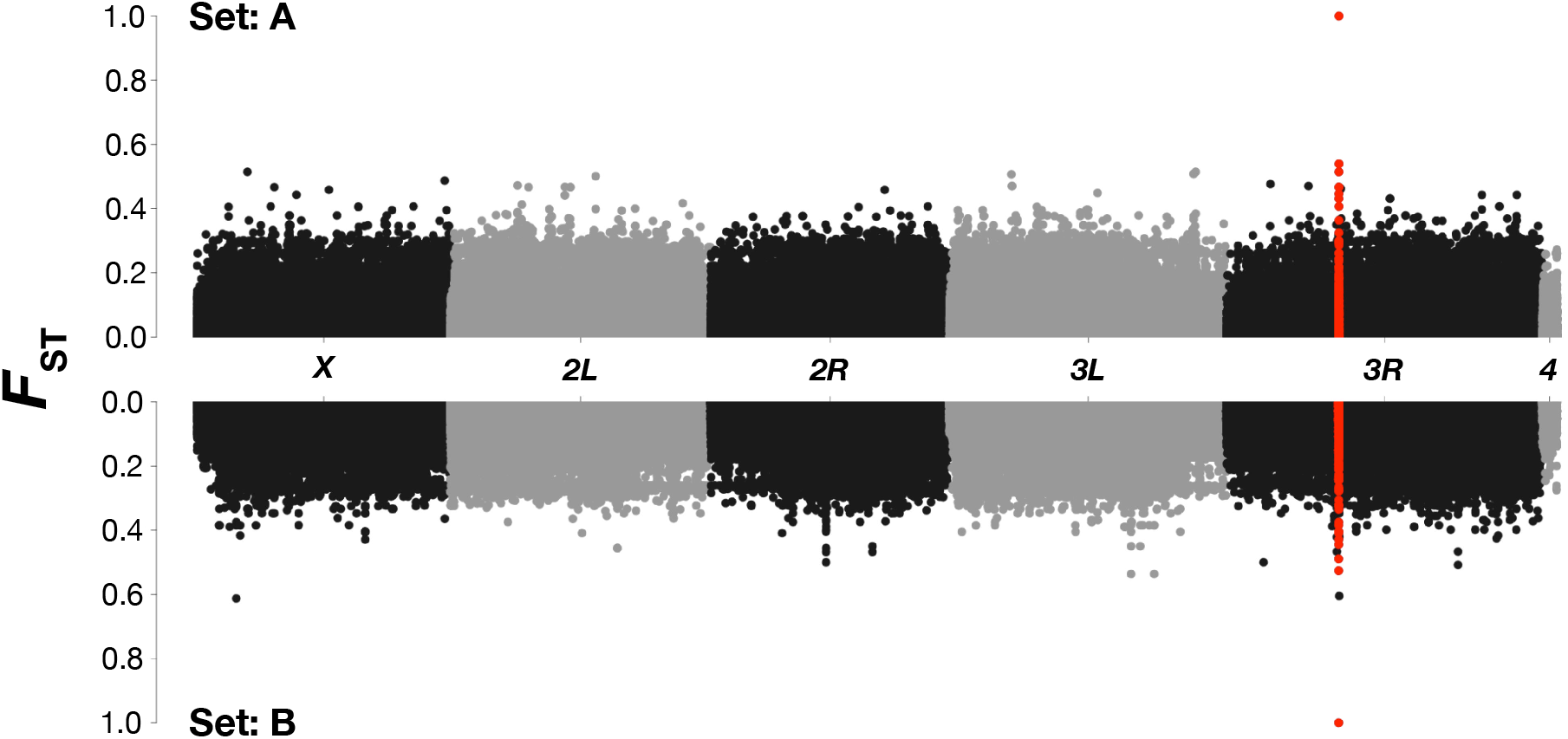
*F*_ST_ Manhattan plots for the biological replicates A and B, constructed from independent sets of inbred lines from the DGRP panel. All SNPs associated with *foxo* are highlighted in red. These analyses show that only the two focal SNPs are fixed for alternative alleles in the low- and high-latitude population cages of sets A and B.

### 3.2 Clinal variation in natural populations

*Body size*. In the six sampled natural populations, thorax length was highly distinct among populations (Table 1), and these patterns of differentiation exhibited a positive association with latitude (Figure 4C). Wing area exhibited qualitatively identical patterns, as did the ratio of wing area to thorax length (Table 1, Figure 4D). Sexes were analyzed separately due to dimorphism and the potential for differential allometry, but exhibited highly similar patterns of size variation among populations and as a function of latitude. As expected, these results are consistent with previous associations between latitude and body size in *D. melanogaster* (Coyne and Beecham 1987; James et al. 1995; de Jong and Bochdanovits 2003).

**Figure 4.**
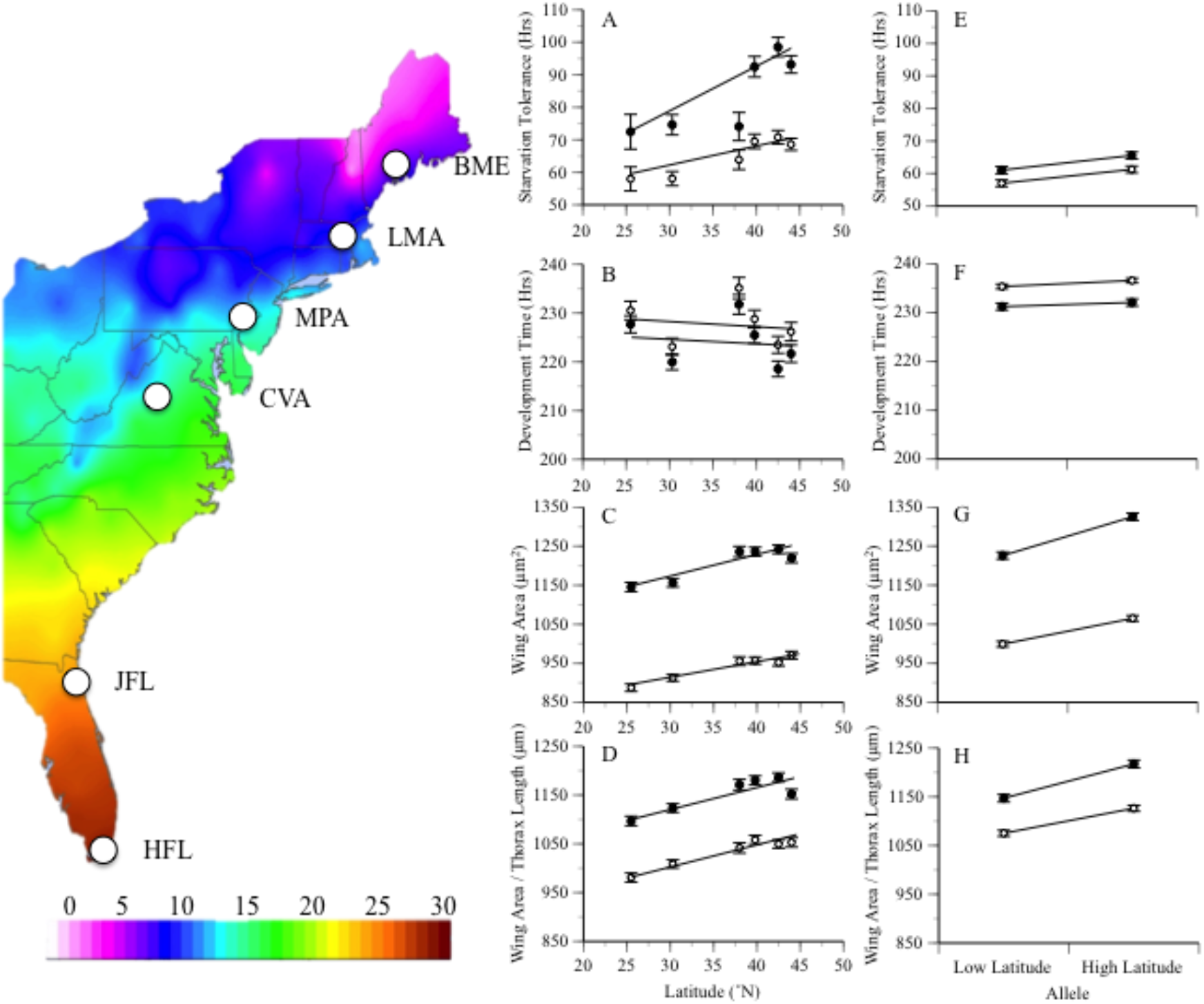
Phenotypic analysis of natural populations collected across the latitudinal gradient in the eastern U.S. (A-D) and the homozygous high- and low-latitude foxo genotypes (E-H). In all panels, females are depicted by filled symbols and males by open symbols. Starvation tolerance increases with increasing latitude (A); similarly, the high-latitude *foxo* allele is associated with increased starvation resistance (E). Development time does not vary predictably with latitude (B), and is also equivalent between *foxo* alleles (F). Wing area (C) and the ratio of wing area to thorax length (D) exhibit a positive latitudinal cline in the sampled populations; these patterns of size variation in the natural populations are mirrored in both magnitude and direction by the observed differences in size parameters between the low and high-latitude foxo alleles (G, H).

**Table 1.**
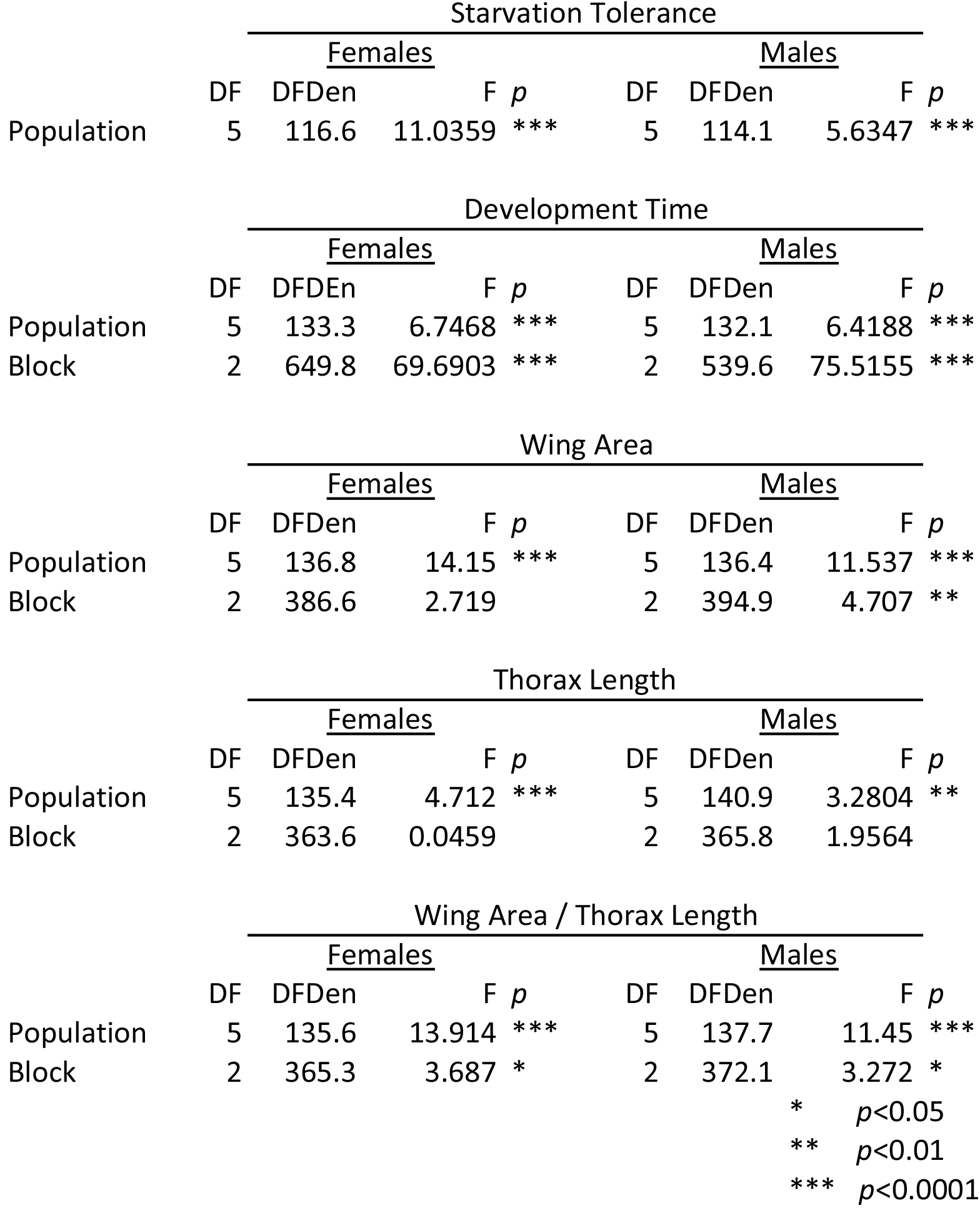
Analyses of variance for the assayed phenotypic traits among natural populations.

*Starvation resistance*. As with body size, starvation tolerance was highly variable among isofemale lines within populations yet exhibited a robust association with geography for both sexes (Table 1, Figure 3A). The increase in starvation tolerance as a function of increasing latitude appears more pronounced for females than males, although patterns are qualitatively identical and no heterogeneity of slopes was detected (analysis not shown). The patterns of increasing tolerance with increasing latitude is consistent with other aspects of stress tolerance in North American populations (Schmidt and Paaby 2008), but opposite to what has been observed on the Indian subcontinent (e.g., Karan et al. 1998). Starvation tolerance does not appear to vary predictably with latitude in other assayed geographic regions (Robinson et al. 2000; Hoffmann et al. 2001).

*Development time*. In contrast to the patterns observed for size and starvation tolerance, development time did not vary predictably with latitude. For both males and females, significant variation was observed among lines and among populations but there was no association with latitudinal origin (Table 1, Figure 4B). We did, however, observe distinct patterns of development time among the replicate experimental blocks, despite controlling for density and culture conditions. This suggests that the trait is affected by additional environmental variables, measurement or other experimental error, or a combination of the two. It should be noted, however, that in examination of the experimental blocks individually, no significant association between development time and latitudinal origin of the population was observed. This is in contrast to patterns of seasonal variation, which demonstrate predictable change in development time as a function of time of collection (Behrman et al. 2015). All raw data for the phenotyping of these natural collections are available (Dryad Accession: XXXXXXX).

### 3.3 Phenotypic differentiation between *foxo* alleles

*Body size*. Thorax length and wing area were both highly distinct between the high and low-latitude *foxo* genotypes (Table 2). Significant heterogeneity was present between the replicate sets **A** and **B**, as expected based on distinct composition of founding inbred lines, as well as among experimental replicates that were cultured independently since cage initiation. Despite these two sources of cage effects, the differences between *foxo* genotypes were consistent with expectations based on geography: the genotypes homozygous for the high-latitude *foxo* allele were significantly larger than genotypes homozygous for the low-latitude allele (Figure 4). The ratio of wing area to thorax length did not show any further differentiation between genotypes, but as with the two traits independently, demonstrated significant and predictable differences between the high and low-latitude *foxo* genotypes (Figure 4H). Furthermore, the observed differences between *foxo* genotypes are strikingly similar in effect size to the trait differences observed between the populations sampled from the latitudinal extremes (Figure 4 C,D vs. Figure 4 G,H).

**Table 2.**
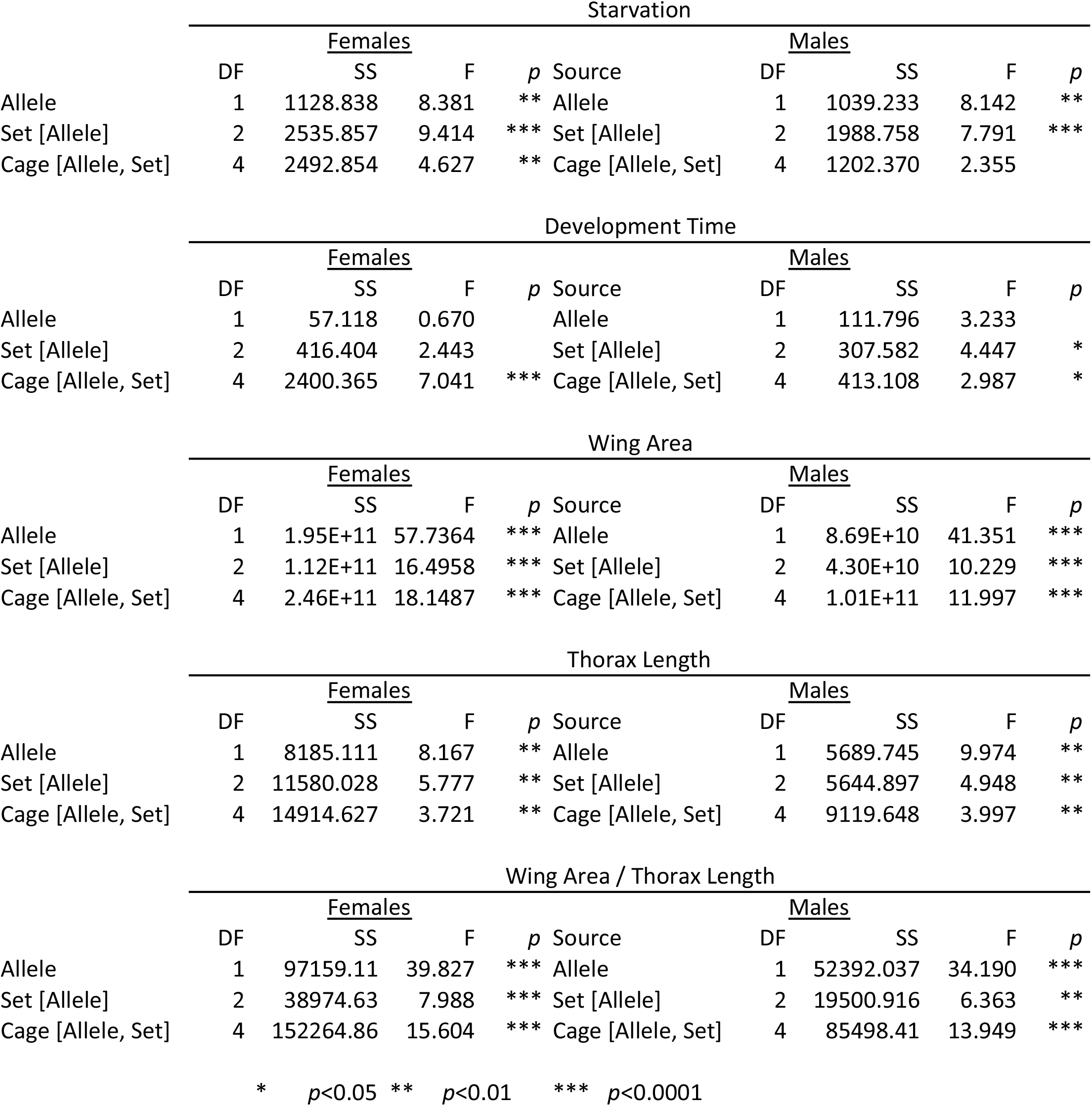
Analyses of variation for the effects of the high and low latitude *foxo* alleles on phenotype.

| Starvation |  |  |  |  |  |  |  |  |  |
| --- | --- | --- | --- | --- | --- | --- | --- | --- | --- |
|  | Females |  |  |  | Source | Males |  |  |  |
|  | DF | SS | F | *p* |  | DF | SS | F | *p* |
| Allele | 1 | 1128.838 | 8.381 | ** | Allele | 1 | 1039.233 | 8.142 | ** |
| Set [Allele] | 2 | 2535.857 | 9.414 | *** | Set [Allele] | 2 | 1988.758 | 7.791 | *** |
| Cage [Allele, Set] | 4 | 2492.854 | 4.627 | ** | Cage [Allele, Set] | 4 | 1202.370 | 2.355 |  |
| Development Time |  |  |  |  |  |  |  |  |  |
|  | Females |  |  |  | Source | Males |  |  |  |
|  | DF | SS | F | *p* |  | DF | SS | F | *p* |
| Allele | 1 | 57.118 | 0.670 |  | Allele | 1 | 111.796 | 3.233 |  |
| Set [Allele] | 2 | 416.404 | 2.443 |  | Set [Allele] | 2 | 307.582 | 4.447 | * |
| Cage [Allele, Set] | 4 | 2400.365 | 7.041 | *** | Cage [Allele, Set] | 4 | 413.108 | 2.987 | * |
| Wing Area |  |  |  |  |  |  |  |  |  |
|  | Females |  |  |  | Source | Males |  |  |  |
|  | DF | SS | F | *p* |  | DF | SS | F | *p* |
| Allele | 1 | 1.95E+11 | 57.7364 | *** | Allele | 1 | 8.69E+10 | 41.351 | *** |
| Set [Allele] | 2 | 1.12E+11 | 16.4958 | *** | Set [Allele] | 2 | 4.30E+10 | 10.229 | *** |
| Cage [Allele, Set] | 4 | 2.46E+11 | 18.1487 | *** | Cage [Allele, Set] | 4 | 1.01E+11 | 11.997 | *** |
| Thorax Length |  |  |  |  |  |  |  |  |  |
|  | Females |  |  |  | Source | Males |  |  |  |
|  | DF | SS | F | *p* |  | DF | SS | F | *p* |
| Allele | 1 | 8185.111 | 8.167 | ** | Allele | 1 | 5689.745 | 9.974 | ** |
| Set [Allele] | 2 | 11580.028 | 5.777 | ** | Set [Allele] | 2 | 5644.897 | 4.948 | ** |
| Cage [Allele, Set] | 4 | 14914.627 | 3.721 | ** | Cage [Allele, Set] | 4 | 9119.648 | 3.997 | ** |
| Wing Area / Thorax Length |  |  |  |  |  |  |  |  |  |
|  | Females |  |  |  | Source | Males |  |  |  |
|  | DF | SS | F | *p* |  | DF | SS | F | *p* |
| Allele | 1 | 97159.11 | 39.827 | *** | Allele | 1 | 52392.037 | 34.190 | *** |
| Set [Allele] | 2 | 38974.63 | 7.988 | *** | Set [Allele] | 2 | 19500.916 | 6.363 | ** |
| Cage [Allele, Set] | 4 | 152264.86 | 15.604 | *** | Cage [Allele, Set] | 4 | 85498.41 | 13.949 | *** |

\* $p < 0.05$ \*\* $p < 0.01$ \*\*\* $p < 0.0001$

*Starvation resistance*. Similar to the observed differences in various metrics for body size, starvation resistance was also distinct between the *foxo* genotypes and varied predictably with geography (Table 2, Figure 4). For both males and females, the genotype homozygous for the high-latitude *foxo* allele was associated with increased starvation tolerance, which may be associated with effects of the alleles on body size and lipid content (e.g., Chippindale et al. 1996). A significant amount of variance in starvation tolerance was also associated with cage effects for both sets of inbred lines and culture replicate (Table 2). The effect size associated with *foxo* genotype was approximately 10%, again similar to what was observed for associated differences in body size. However, unlike the patterns observed for size related traits, the differences between *foxo* genotypes appear to explain a small amount of the variance in this trait among natural populations across the sampled geographic range in the eastern U.S. (Figure 4A, H). These distinct patterns suggest that differences in starvation tolerance are not determined exclusively by differences in size.

*Development time*. Development time varied significantly across sets of lines and replicates; in contrast to the other traits measured, development time was not distinct between the high- and low-latitude *foxo* alleles (Table 2, Figure 4). Overall these results suggest that there is no functional significance associated with these naturally occurring *foxo* alleles with respect to development time. This is somewhat congruent with data on Australian clines, where the relationship between body size and development time is inconsistent across latitude (e.g., James et al 1995, 1997). Development time was the most variable of the traits studied here, both with respect to experimental replication and variation among natural as well as reconstituted outbred populations.

## 4. DISCUSSION

In *D. melanogaster*, there is abundant evidence for local adaptation. Natural populations exhibit rapid and predictable responses to environmental parameters that vary with season, both in terms of phenotypic (Schmidt and Conde 2006; Behrman et al. 2015; Rajpurohit et al. 2017; Behrman et al. 2018a; Behrman et al. 2018b; Rajpurohit et al. 2018) and allele frequency (Cogni et al. 2014; Bergland et al. 2014; Behrman et al. 2018a; Machado et al. 2018) change. Similarly, many fitness-associated traits have been shown to vary predictably with latitude, often in parallel across independent gradients (e.g., Oakeshott et al. 1982; Paaby et al. 2010; Yang and Edery 2018). Latitudinal allele frequency clines at candidate loci (e.g., Schmidt et al. 2000; Betancourt et al. 2001; Sezgin et al. 2005; Paaby et al. 2010; Cogni et al. 2017) are now placed in a genomic context in which tens of thousands of SNPs are known to be clinal (e.g., Kolackzowski et al. 2011; Fabian et al. 2012; Bergland et al. 2016; Kapun et al. 2016). While it is clear that some allele frequency clines may be generated by demography (Kao et al. 2015; Bergland et al. 2016), it is also clear that at least some of the observed clines may be generated by spatially varying selection (Schmidt et al. 2008; Svetec et al. 2016). Two of the associated, major questions are: 1) how many, or what proportion of, allele frequency clines reflect spatially varying selection and thus local adaptation; and 2) how are allele frequency and phenotypic clines integrated? Alleles exist in a genomic context, and complex traits are similarly correlated as well as affected by epistasis (Mackay 2014); it is extremely unlikely that all allele and phenotypic clines are independent and reflect selection on single variants or traits. However, the effect size of individual “adaptive polymorphisms” and the architecture of local adaptation are, arguably, not well resolved. There is a paucity of detailed, mechanistic and comprehensive investigations as to the functional significance of segregating polymorphisms in natural populations. It is clearly infeasible, as well as misguided, to assess the functional impact of all polymorphisms across the genome. However, we suggest that multiple, intersecting methodologies (e.g., direct mapping, expression analyses, mutant analysis, patterns of variation in natural populations) can be used to identify a subset of variants that may be examined for functional significance. Ideally, investigation of a sufficient number of outliers could generate an empirical distribution of genic or allelic effect sizes for specific fitness-associated traits. Such investigations are essential in resolving the genetic architecture and dynamics of local adaptation.

The *foxo* alleles we examine here are an example of a robust candidate suitable for functional analysis. Genetic manipulations of the *foxo* gene have revealed pronounced effects on lifespan, multiple aspects of stress tolerance including starvation resistance, and growth phenotype (Jünger et al., 2003; Puig, Marr, Ruhf, & Tjian, 2003; Kramer, Davidge, Lockyer, & Staveley, 2003; Hwangbo, Gersham, Tu, Palmer, & Tatar, 2004; Giannakou et al. 2004; Kramer, Slade, Staveley, 2008; Slack, Giannakou, Foley, Goss, & Partridge, 2011). These traits vary with latitude in *D. melanogaster* (Coyne & Beecham 1987; James et al. 1995; Schmidt et al. 2005); thus, the *foxo* gene is a logical candidate for determining variation for these traits in natural populations (de Jong and Bochdonavits 2003).

However, the effects of natural variation have not been previously addressed. While loss of function mutants are highly pleiotropic and have pronounced effects on life history phenotypes, and the IIS/TOR pathway is in general extremely pleiotropic, polymorphisms segregating in natural populations need not affect variance for all traits related to gene function (e.g., Stern 2011).

Our results demonstrate that allelic variation at *foxo* affects body size and starvation tolerance. We further show that these measures of size, thorax length and wing area, as well as starvation tolerance, vary predictably with latitude in natural populations of *D. melanogaster*. Furthermore, the high-latitude *foxo* allele is associated with larger size and greater starvation resistance, whereas flies homozygous for the low-latitude *foxo* allele are smaller and less tolerant. Thus, the allelic effects also parallel patterns of variation in natural populations. This is distinct from the countergradient patterns that have been previously observed for allelic variants at the *Drosophila insulin receptor* (*InR*) (Paaby et al. 2014). The effect size of the *foxo* alleles is seemingly large, particularly for body size: the difference between high and low-latitude *foxo* alleles is approximately the same as the observed size difference between flies sampled from Florida and Maine. This suggests that allelic variation at *foxo* is a major contributor to variance in body size in these populations. It remains to be determined whether *foxo* is a major effect locus for size in other taxa.

Durmaz et al. (2018) demonstrate that these focal *foxo* variants also have significant effects on viability, fat catabolism, and FOXO activity, as indicated by differences in transcript abundance of a FOXO target (*InR*). These investigations were done after an additional four generations of recombination, and parallel differences across studies were observed for different measures of body size: wing area, thorax length, the ratio of wing area:length, and femur length were all greater in the populations homozygous for the high-latitude allele than in the recombinant populations homozygous for the low-latitude allele. Differences between the high and low-latitude *foxo* variants were largely consistent across two assayed temperatures and two diets, although genotype by environment interactions were observed for both starvation tolerance and lipid metabolism (Durmaz et al. 2018). While our results for *foxo* allelic effects are consistent with those of Durmaz et al. (2018) for measures of size, patterns of starvation tolerance for the high and low-latitude alleles are opposite: in our data, populations with the high-latitude *foxo* allele were more starvation tolerant, whereas in Durmaz et al. (2018) the low-latitude allele was associated with increased tolerance. The discrepancy may be due to methodological disparity, as the two assays are distinct; our method is associated with faster mortality and may involve a greater degree of desiccation stress than the agar method. Desiccation tolerance does vary with latitude in North American populations (Rajpurohit et al. 2018), although the effects of *foxo* on desiccation tolerance are unknown. Alternatively, the one observed difference across the two studies may be associated with laboratory-specific microbiota, which is known to vary across labs and culture conditions (Staubach et al. 2012) and has pronounced effects on *D. melanogaster* life histories (Walters et al., in review).

Despite the parallels we observed between the assayed *foxo* variants and the patterns in natural populations, we cannot conclude that it is these two focal SNPs (positions) that themselves cause the observed differences in starvation tolerance and size, or directly contribute to variance for these traits in natural populations. The linkage disequilibrium present in the founding inbred lines (DGRP) would decay to some extent by the eight generations of outcrossing among founder lines, but remains pronounced; thus, without further characterization these SNPs are interpreted as markers for the functionally significant allelic variation segregating at this locus. Gene editing or similar techniques, in which the focal sites are manipulated in multiple common genetic backgrounds, would be essential in directly examining causality. Such investigations are the focus of future work.

## 5. CONCLUSION

We find that the marker allele at *foxo* has predictable effects on body size and starvation tolerance; no effects of *foxo* alleles on development time were observed. We also show that both starvation tolerance and body size exhibit pronounced latitudinal clines in six sampled natural populations, whereas development time exhibited no association with latitude. The assayed alleles at *foxo* explain a small amount of the variance among natural populations for starvation tolerance, but appear to be a major factor in the determination of variance in size. Our results suggest a distinct genetic architecture for correlated fitness traits, and that allelic variation at the *foxo* locus underlies, in part, patterns of local adaptation in natural populations of this genetic model.

## ACKNOWLEDGEMENTS

We thank members of the Flatt and Schmidt labs for their gracious assistance in all aspects of this work. Financial support was provided by the National Institutes of Health (R01GM100366 to P.S.), the U.S. National Science Foundation (DEB0921307 to P.S.), the Austrian Science Foundation (FWF P21498-B11 to T.F.), and the Swiss National Science Foundation (PP00P3_133641 to T.F.).

## DATA ACCESSIBILITY

The raw phenotypic data are available from Dryad at: XXXXX.

## AUTHOR CONTRIBUTIONS

P.S. and T.F. conceived the project. D.F. and M.K. identified the *foxo* SNPs and performed genomic analyses. P.S., S.R., E.D. and T.F. designed the experiments. N.B. and S.R. established populations and performed the experiments. P.S., N.B., E.D., M.K., S.R. and T.F. analyzed the data. N.B., T.F. and P.S. wrote the paper with input from the other authors.

## LITERATURE CITED

Arnegard, M. E., McGee, M. D., Matthews, B., Marchinko, K. B., Conte, G. L., Kabir, S., … Schluter, D. (2014). Genetics of ecological divergence during speciation. Nature, 511(7509), 307–311. http://doi.org/10.1038/nature13301

Ashton KG. (2002). Patterns of within-species body size variation of birds: strong evidence for Bergmann’s rule. Global Ecol. Biogeogr. 11:505–523.

Azevedo, R.B.R., James, A.C., McCabe, J., & Partridge, L. (1998). Latitudinal variation of wing: thorax size ratio and wing aspect ratio in *Drosophila melanogaster*. Evolution, 52, 1353–1362. https://doi.org/10.1111/j.1558-5646.1998.tb02017.x.

Barton N.H., A.M. Etheridge, A. Veber. (2017). The infinitesimal model: Definition, derivation, and implications. Theoretical Population Biology 118 (2017) 50–73.

Behrman, E. L., Watson, S. S., O’brien, K. R., Heschel, M. S., & Schmidt, P.S. (2015). Seasonal variation in life history traits in two *Drosophila* species. Journal of Evolutionary Biology, 28(9), 1691–1704. http://doi.org/10.1111/jeb.12690

Behrman, E. L., Howick, V. M., Kapun, M., Staubach, F., Bergland, A. O., Petrov, D. A.,. Schmidt, P. S. (2018). Rapid seasonal evolution in innate immunity of wild *Drosophila melanogaster*. Proceedings of the Royal Society of London B, 285, 20172599. https://doi.org/10.1098/rspb.2017.2599

Berhman, E.L., Kawecki, T.J., & Schmidt, P. (2018). Natural variation in couch potato mediates rapid evolution of learning and reproduction in natural populations of Drosophila melanogaster. doi: https://doi.org/10.1101/288696

Bergland, A. O., Behrman, E. L., O’Brien, K. R., Schmidt, P. S., & Petrov, D. A. (2014). Genomic evidence of rapid and stable adaptive oscillations over seasonal time scales in *Drosophila*. PLoS Genetics, 10, e1004775. https://doi.org/10.1371/journal.pgen.1004775

Bergland, A. O., Tobler, R., González, J., Schmidt, P., & Petrov, D. (2016). Secondary contact and local adaptation contribute to genome-wide patterns of clinal variation *in Drosophila melanogaster*. Molecular Ecology, 25, 1157–1174. https://doi.org/10.1111/mec.13455

Blanckenhorn W. U. 2000. The evolution of body size: What keeps organisms small? Q. Rev. Biol. 75, 385–407.

Blanckenhorn, W.U. and M. Demont 2004. Bergmann and converse bergmann latitudinal clines in arthropods: two ends of a continuum? Integr Comp Biol. 44(6):413–24.

Bonnet T, Wandeler P, Camenisch G, Postma E (2017) Bigger Is Fitter? Quantitative Genetic Decomposition of Selection Reveals an Adaptive Evolutionary Decline of Body Mass in a Wild Rodent Population. PLoS Biol 15(1): e1002592. https://doi.org/10.1371/journal.pbio.1002592

Boyle EA, Yi Li, JK Pritchard. (2017). An expanded view of complex traits: from polygenic to omnigenic. Cell 169 (7), 1177–1186.

Brown JH, Marquet PA, Taper ML. (1993). Evolution of body size: consequences of an energetic definition of fitness. Am Nat.142(4):573–84. doi: 10.1086/285558.

Chakraborty, M., and J. D. Fry. (2016). Evidence that environmental heterogeneity maintains a detoxifying enzyme polymorphism in *Drosophila melanogaster*. Current Biology 26:219–223.

Cheviron, Z. A., G. C. Bachman, A. D. Connaty, G. B. McClelland, and J. F. Storz. (2012). Regulatory changes contribute to the adaptive enhancement of thermogenic capacity in high-altitude deer mice. Proceedings of the National Academy of Sciences USA, 109: 8635–8640.

Chippindale A. K., Chu T. J. F. and Rose M. R. 1996. Complex trade-offs and the evolution of starvation resistance in *Drosophila melanogaster*. Evolution 50, 753–766.

Clancy DJ, Gems D, Harshman LG, Oldham S, Stocker H, Hafen E, Leevers SJ, Partridge L. 2001. Extension of life-span by loss of CHICO, a Drosophila insulin receptor substrate protein. Science 2001; 292:41–3.

Clemente F, Vogl C (2012) Unconstrained evolution in short introns? – An analysis of genome-wide polymorphism and divergence data from *Drosophila*. Journal of Evolutionary Biology, 25, 1975–1990.

Cogni R, Kuczynski C, Koury S, Lavington E, Behrman EL, O’Brien KR, Schmidt PS, Eanes WF. (2014).The intensity of selection acting on the couch potato gene-spatial-temporal variation in a diapause cline. Evolution 68(2):538–48. doi: 10.1111/evo.12291.

Cogni, R., Kuczynski, K., Koury, S., Lavington, E., Behrman, E. L., O’Brien, K. R., … Eanes, W. F. (2017). On the Long-term Stability of Clines in Some Metabolic Genes in *Drosophila melanogaster*. Scientific Reports, 7, 42766. http://doi.org/10.1038/srep42766

Colombani J, Bianchini L, Layalle S, Pondeville E, Dauphin-Villemant C, Antoniewski C, Carré C, Noselli S, Léopold P. (2005). Antagonistic actions of ecdysone and insulins determine final size in Drosophila. Science 310(5748):667–70.

Colosimo PF, Hosemann KE, Balabhadra S, Villarreal G Jr, Dickson M, Grimwood J, Schmutz J, Myers RM, Schluter D, Kingsley DM. (2005). Widespread parallel evolution in sticklebacks by repeated fixation of Ectodysplasin alleles. Science. 307(5717):1928–33.

Corbett-Detig RB, Cardeno C, Langley CH (2012) Sequence-based detection and breakpoint assembly of polymorphic inversions. Genetics, 192, 131–137.

Comeault AA, Flaxman SM, Riesch R, Curran E, Soria-Carrasco V, Gompert Z, Farkas TE, Muschick M, Parchman TL, Schwander T, Slate J, Nosil P. (2015). Selection on a genetic polymorphism counteracts ecological speciation in a stick insect. Curr Biol. 25(15):1975–81. doi: 10.1016/j.cub.2015.05.058.

Coyne J. A. and Beecham E. 1987. Heritability of two morphological characters within and among natural populations of Drosophila melanogaster. Genetics 117: 727–737.

David J. R. and Capy P. 1988. Genetic variation of Drosophila melanogaster natural populations. Trends Genet. 4, 106–111.

de Jong G. and Bochdanovits Z. 2003 Latitudinal clines in Drosophila melanogaster: body size, allozyme frequencies, inversion frequencies, and the insulin-signalling pathway. J. Genet. 82, 207–223

Djawdan M., Chippindale A. K., Rose M. R. and Bradley T. J. 1998. Metabolic reserves and evolved stress resistance in Drosophila melanogaster. Physiol. Zool. 71, 584–594.

Durmaz, E., Rajpurohit, S., Betancourt, N., Fabian, D.K., Kapun, M., Schmidt, P., & Flatt, T. (2018). A clinal polymorphism in the insulin signaling transcription factor foxo contributes to life history adaptation in Drosophila. Molecular Ecology, in review.

Fabian DK, Kapun M, Nolte V, Kofler R, Schmidt PS, Schlotterer C, and Flatt T. 2012. Genome-wide patterns of latitudinal differentiation among populations of Drosophila melanogaster from North America. Molecular Ecology (2012). Blackwell Publishing Ltd.

Fabian, D. K., Lack, J. B., Mathur, V., Schlötterer, C., Schmidt, P. S., Pool, J. E.,. Flatt, T. (2015). Spatially varying selection shapes life history clines among populations of *Drosophila melanogaster* from sub-Saharan Africa. Journal of Evolutionary Biology, 28, 826–840. https://doi.org/10.1111/jeb.12607

Fielenbach, N., & Antebi, A. (2008). C. elegans dauer formation and the molecular basis of plasticity. Genes & Development, 22, 2149–2165. https://doi.org/10.1101/gad.1701508

Garofalo R. S. 2002. Genetic analysis of insulin signaling in Drosophila. Trends Endocrinol. Metab. 13, 156–162.

Giannakou ME, Goss M, Juenger MA, Hafen E, Leevers SJ, Partridge L. 2004. Long-lived Drosophila with overexpressed dFOXO in adult fat body. Science 2004; 305:361.

Giannakou ME, Partridge L. 2007. Role of insulin-like signalling in Drosophila lifespan. Trends in Biochemical Science, 32, 180–188.

Gilchrist, A.S., Azevedo, R.B.R., Partridge, L., & O’Higgins, P. (2000). Adaptation and constraint in the evolution of *Drosophila melanogaster* wing shape. Evolution & Development, 2, 114–124. https://doi.org/10.1046/j.1525-142x.2000.00041.x

Harshman L. G., Hoffmann A. A. and Clark A. G. 1999. Selection for starvation resistance in Drosophila melanogaster: physiological correlates, enzyme activities and multiple stress responses. J. Evol. Biol. 12, 370–379.

Hoffmann A. A., Hallas R., Sinclair C. and Mitrovski P. 2001. Levels of variation in stress resistance in Drosophila among strains, local populations and geographic regions: Patterns for desiccation, starvation, cold resistance, and associated traits. Evolution 55, 1621–1630.

Huey, R.B., G.W. Gilchrist, M.L. Carson, D. Berrigan and L. Serra. (2000). Rapid evolution of a geographic cline in size in an introduced fly. Science 287: 308–309.

Hwangbo, D. S., Gersham, B., Tu, M.-P., Palmer, M., & Tatar, M. (2004). *Drosophila* dFOXO controls lifespan and regulates insulin signalling in brain and fat body. Nature, 429, 562–566. https://doi.org/10.1038/nature02549

James A. C. and Partridge L. 1995. Thermal evolution of rate of larval development in Drosophila melanogaster in laboratory and field populations. J. Evol. Biol. 8, 315–330.

James A. C., Azevedo R. B. R. and Partridge L. 1995. Cellular basis and developmental timing in a size cline of Drosophila melanogaster. Genetics 140, 659–666.

James A. C., Azevedo R. B. R. and Partridge L. 1997. Genetic and environmental responses to temperature of Drosophila melanogaster from a latitudinal cline. Genetics 146, 881–890.

Jones MR, Mills LS, Alves PC, Callahan CM, Alves JM, Lafferty DJR, Jiggins FM, Jensen JD, Melo-Ferreira J, Good JM. (2018). Adaptive introgression underlies polymorphic seasonal camouflage in snowshoe hares. Science. 360(6395):1355–1358. doi: 10.1126/science.aar5273.

Kao, J. Y., Zubair, A., Salomon, M. P., Nuzhdin, S. V., & Campo, D. (2015). Population genomic analysis uncovers African and European admixture in *Drosophila melanogaster* populations from the south-eastern United States and Caribbean Islands. Molecular Ecology, 24, 1499–1509. https://doi.org/10.1111/mec.13137

Kapun M, Fabian DK, Goudet J, Flatt T (2016) Genomic Evidence for Adaptive Inversion Clines in *Drosophila melanogaster*. Molecular Biology and Evolution, 33, 1317–1336.

Karan D., Dahiya N., Munjal A. K., Gibert P., Moreteau B., Parkash R. and David J. R. 1998. Desiccation and starvation tolerance of adult Drosophila: Opposite latitudinal clines in natural populations of three different species. Evolution 52, 825–831.

Kolaczkowski, B., Kern, A. D., Holloway, A. K., & Begun, D. J. (2011). Genomic differentiation between temperate and tropical Australian populations of *Drosophila melanogaster*. Genetics, 187, 245–260. https://doi.org/10.1534/genetics.110.123059

Kramer, J. M., Davidge, J. T., Lockyer, J. M., & Staveley, B. E. (2003). Expression of *Drosophila* FOXO regulates growth and can phenocopy starvation. BMC Developmental Biology, 3, 5. https://dx.doi.org/10.1186%2F1471-213X-3-5

Kramer, J. M., Slade, J. D., & Staveley, B. E. (2008). *foxo* is required for resistance to amino acid starvation in Drosophila. Genome, 51, 668–672. https://doi.org/10.1139/G08-047

Lack JB, Monette MJ, Johanning EJ, Sprengelmeyer QD, Pool JE (2016) Decanalization of wing development accompanied the evolution of large wings in high altitude *Drosophila*. Proc Natl Acad Sci USA 113:1014–1019.

Langley CH, Stevens K, Cardeno C et al. (2012) Genomic variation in natural populations of *Drosophila melanogaster*. Genetics, 192, 533–598.

Laurie, CC and LF Stam. (1988). Quantitative analysis of RNA produced by Slow and Fast alleles of *Drosophila melanogaster*. Proc. Natl. Acad. Sci. USA 85: 5161–5165.

Lamichhaney S, Han F, Berglund J, Wang C, Almen MS, Webster MT, et al. (2016). A beak size locus in Darwins finches facilitated character displacement during a drought. Science. 352(6284):470–474. doi: 10.1126/science.aad8786 PMID: 27102486

Lewontin, R. C. (1974). The Genetic Basis of Evolutionary Change. New York, USA: Columbia University Press.

Libina, N., Berman, J. R., & Kenyon, C. (2003). Tissue-specific activities of *C. elegans* DAF-16 in the regulation of lifespan. Cell, 115, 489–502. https://doi.org/10.1016/S0092-8674(03)00889-4

Machado, H. E., Bergland, A. O., Taylor, R., Tilk, S., Behrman, E. L., Dyer, K., … Petrov, D. (2018). Broad geographic sampling reveals predictable and pervasive seasonal adaptation in *Drosophila*. bioRxiv 337543. https://doi.org/10.1101/337543

Mackay, T. F. C. (2014). Epistasis and Quantitative Traits: Using Model Organisms to Study Gene-Gene Interactions. Nature Reviews. Genetics, 15(1), 22–33. http://doi.org/10.1038/nrg3627

Mackay TF, Richards S, Stone EA, Barbadilla A, Ayroles JF, Zhu D, Casillas S, Han Y, Magwire MM, Cridland JM, Richardson MF, Anholt RRH, Barrón M, Bess C, Blankenburg KP, Carbone MA, Castellano D, Chaboub L, Duncan L, Harris Z, Javaid M, Jayaseelan JC, Jhangiani SN, Jordan KW, Lara F. 2012. The Drosophila melanogaster Genetic Reference Panel. Nature 482, 173–178.

Mattila, J., Bremer, A., Ahonen, L., Kostiainen, R., & Puig, O. (2009). *Drosophila* FoxO Regulates Organism Size and Stress Resistance through an Adenylate Cyclase. Molecular and Cellular Biology, 29, 5357–5365. https://doi.org/10.1128/MCB.00302-09

Manceau M, Domingues VS, Mallarino R, Hoekstra HE. The developmental role of Agouti in color pattern evolution. Science 2011;331:1062–5.

McKown AD, Klápšt? J, Guy RD, Geraldes A, Porth I, Hannemann J, Friedmann M, Muchero W, Tuskan GA, Ehlting J, Cronk QC, El-Kassaby YA, Mansfield SD, Douglas CJ. (2014). Genome-wide association implicates numerous genes underlying ecological trait variation in natural populations of *Populus trichocarpa*. New Phytol. 203(2):535–53. doi: 10.1111/nph.12815.

Noach E. J. K., de Jong G. and Scharloo W. 1996. Phenotypic plasticity in morphological traits in two populations of Drosophila melanogaster. J. Evol. Biol. 9, 831–844.

Oakeshott JG, Gibson JB, Anderson PR, Knibb WR, Anderson DG, Chambers GK. (1982). Alcohol dehydrogenase and glycerol-3-phosphate dehydrogenase clines in Drosophila melanogaster on different continents. Evolution 36(1):86–96. doi:10.1111/j.1558-5646.1982.tb05013.x.

Oldham S. and Hafen E. 2003. Insulin/IGF and target of rapa-mycin signaling: a TOR de force in growth control. Trends Cell Biol. 13, 79–85.

Paaby AB, Schmidt PS. 2008. Functional significance of allelic variation at methuselah, an aging gene in Drosophila. PLoS ONE, 3, e1987.

Paaby, A. B., & Schmidt, P. S. (2009). Dissecting the genetics of longevity in *Drosophila melanogaster*. Fly, 3, 1–10. https://doi.org/10.4161/fly.3.1.7771

Paaby, A. B., Bergland, A. O., Behrman, E. L., & Schmidt, P. S. (2014). A highly pleiotropic amino acid polymorphism in the *Drosophila* insulin receptor contributes to life-history adaptation. Evolution, 68, 3395–3409. https://doi.org/10.1111/evo.12546

Paaby, A. B., Blacket, M. J., Hoffmann, A. A., & Schmidt, P. S. (2010). Identification of a candidate adaptive polymorphism for *Drosophila* life history by parallel independent clines on two continents. Molecular Ecology, 19, 760–774. https://doi.org/10.1111/j.1365-294X.2009.04508.x

Partridge L, Coyne JA. (1997). Bergmann’s rule in ectotherms: is it adaptive? Evolution. 51:632–635.

Parsch J, Novozhilov S, Saminadin-Peter SS, Wong KM, Andolfatto P (2010) On the utility of short intron sequences as a reference for the detection of positive and negative selection in *Drosophila*. Molecular Biology and Evolution, 27, 1226–1234.

R Development Core Team (2009) R: A Language and Environment for Statistical Computing. R-project.org.

Rajpurohit, S., Hanus, R., Vrkoslav, V., Behrman, E. L., Bergland, A. O., Petrov, D., … Schmidt, P. S. (2017). Adaptive dynamics of cuticular hydrocarbons in *Drosophila*. Journal of Evolutionary Biology, 30(1), 66–80. http://doi.org/10.1111/jeb.12988

Rajpurohit S, Gefen E, Bergland AO, Petrov DA, Gibbs AG, Schmidt PS. (2018). Spatiotemporal dynamics and genome-wide association analysis of desiccation tolerance in *Drosophila melanogaster*. Mol Ecol. 27(17):3525–3540. doi: 10.1111/mec.14814.

Reinhardt, J. A., Kolaczkowski, B., Jones, C. D., Begun, D. J., & Kern, A. D. (2014). Parallel Geographic Variation in *Drosophila melanogaster*. Genetics, 197, 361–373. https://doi.org/10.1534/genetics.114.161463

Robinson S. J. W., Zwaan B. and Partridge L. 2000. Starvation resistance and adult body composition in a latitudinal cline of Drosophila melanogaster. Evolution 54, 1819–1824.

Robinson S. J. W. and Partridge L. 2001. Temperature and cli-nal variation in larval growth efficiency in Drosophila melanogaster. J. Evol. Biol. 14, 14–21.

Rockman, M. V. (2012). The QTN program and the alleles that matter for evolution: all that’s gold does not glitter. Evolution, 66, 1–17. https://doi.org/10.1111/j.1558-5646.2011.01486.x

Roff DA (2007). A centennial celebration for quantitative genetics. Evolution. 61(5):1017–1032. doi: 10.1111/j.1558-5646.2007.00100.x PMID: 17492957

Saucedo L. J. and Edgar B. A. 2002. Why size matters: altering cell size. Curr. Opin. Genet. Dev. 12, 565–571.

Savolainen O, Lascoux M, Merilä J. (2013). Ecological genomics of local adaptation. Nat Rev Genet. 14(11):807–20. doi: 10.1038/nrg3522.

Schmidt, P. S., & Conde, D. R. (2006). Environmental Heterogeneity and the Maintenance of Genetic Variation for Reproductive Diapause in *Drosophila melanogaster*. Evolution, 60, 1602. https://doi.org/10.1111/j.0014-3820.2006.tb00505.x

Schmidt, P. S., & Paaby, A. B. (2008). Reproductive Diapause and Life-History Clines in North American Populations of *Drosophila melanogaster*. Evolution, 62, 1204–1215. https://doi.org/10.1111/j.1558-5646.2008.00351.x

Schmidt, P. S., Duvernell, D. D., & Eanes, W. F. (2000). Adaptive evolution of a candidate gene for aging in *Drosophila*. Proceedings of the National Academy of Sciences, 97, 10861–10865. https://dx.doi.org/10.1073%2Fpnas.190338897

Schmidt, P. S., Matzkin, L., Ippolito, M., & Eanes, W. F. (2005). Geographic Variation in Diapause Incidence, Life-History Traits, and Climatic Adaptation in *Drosophila melanogaster*. Evolution, 59, 1721–1732. https://doi.org/10.1111/j.0014-3820.2005.tb01821.x

Schmidt, P. S., Paaby, A. B., & Heschel, M. S. (2005). Genetic variance for diapause expression and associated life histories in *Drosophila melanogaster*. Evolution, 59, 2616–2625. https://doi.org/10.1111/j.0014-3820.2005.tb00974.x

Schmidt, P. S., Zhu, C-T., Das, J., Batavia, M., Yang, L., & Eanes, W. F. (2008). An amino acid polymorphism in the *couch potato* gene forms the basis for climatic adaptation in *Drosophila melanogaster*. Proceedings of the National Academy of Sciences USA, 105, 16207–16211. https://doi.org/10.1073/pnas.0805485105

Sezgin, E., Duvernell, D. D., Matzkin, L. M., Duan, Y., Zhu, C.-T., Verrelli, B. C., & Eanes, W. F. (2004). Single-Locus Latitudinal Clines and Their Relationship to Temperate Adaptation in Metabolic Genes and Derived Alleles in *Drosophila melanogaster*. Genetics, 168(2), 923–931. http://doi.org/10.1534/genetics.104.027649

Sim C, Kang DS, Kim S, Bai X, and Denlinger D. 2015. Identification of FOXO targets that generate diverse features of the diapause phenotype in the mosquito *Culex pipiens*. PNAS. March 24, 2015 | vol. 112 | no. 12 | 3811–3816

Staubach, F., Baines, J. F., Künzel, S., Bik, E. M., & Petrov, D. A. (2013). Host Species and Environmental Effects on Bacterial Communities Associated with *Drosophila* in the Laboratory and in the Natural Environment. PLoS ONE, 8(8), e70749. http://doi.org/10.1371/journal.pone.0070749

Stern, D. L., (2011). Evolution, development, & the predictable genome. Roberts & Co. Publishers. https://doi.org/10.1086/663915

Stillwell, R.C. (2010). Oikos. 119(9): 1387–1390. doi: 10.1111/j.1600-0706.2010.18670.x

Stillwell RC, Morse GE, Fox CW. (2007). Geographic variation in body size and sexual size dimorphism of a seed-feeding beetle.Am Nat. 170(3):358–69.

Sutter, N. B., Bustamante, C. D., Chase, K., Gray, M. M., Zhao, K., Zhu, L., … Ostrander, E. A. (2007). A Single *IGF1* Allele Is a Major Determinant of Small Size in Dogs. Science (New York, N.Y.), 316(5821), 112–115. http://doi.org/10.1126/science.1137045

Svetec, N., Cridland, J. M., Zhao, L., & Begun, D. J. (2016). The Adaptive Significance of Natural Genetic Variation in the DNA Damage Response of *Drosophila melanogaster*. PLoS Genetics, 12(3), e1005869. http://doi.org/10.1371/journal.pgen.1005869

Tatar M, Kopelman A, Epstein D, Tu M-P, Yin C-M, Garofalo RS. 2001. A mutant Drosophila insulin receptor homolog that extends life-span and impairs neuroendocrine function. Science 2001; 292:107–10

Tatar M, Yin C. (2011). Slow aging during insect reproductive diapause: Why butterflies, grasshoppers and flies are like worms. Exp Gerontol 36(4-6):723–738.

Umina, PA, AR Weeks, MR Kearney, SW McKechnie and AA Hoffmann. (2005). A rapid shift in a classic clinal pattern in Drosophila reflecting climate change. Science 308(5722): 691–693.

van’t Hof, A.E., Campagne, P., Rigden, D.J., Yung, C.J., Lingley, J., Quail, M.A., Hall, N., Darby, A.C., and Saccheri, I.J. (2016). The industrial melanism mutation in British peppered moths is a transposable element. Nature 534, 102–105.

Walters, AW, Matthews, MK, Hughes, RC, Malcolm, J, Rudman, SR, Newell, PD, Douglas, A, Schmidt, P, & Chaston, JM. 2018. The microbiota influences the Drosophila melanogaster life history strategy. PLoS Biology, in review.

Weir BS, Cockerham CC (1984) Estimating *F*-Statistics for the Analysis of Population Structure. Evolution, 38, 1358–1370.

Wellenreuther M, Hansson B. (2016). Detecting Polygenic Evolution: Problems, Pitfalls, and Promises. Trends Genet. 32(3):1–10. doi: 10.1016/j.tig.2015.12.004

Yang, Y. and I. Edery. (2018). Parallel clinal variation in the mid-day siesta of *Drosophila melanogaster* implicates continent-specific targets of natural selection. PloS Genetics. https://doi.org/10.1371/journal.pgen.1007612

Zwaan B. J., Azevedo R. B. R., James A. C., van’t Land J. and Partridge L. 2000. Cellular basis of wing size variation in Drosophila melanogaster: a comparison of latitudinal clines on two continents. Heredity 84, 338–347.

